# Effects of Controlled Dual Growth Factor Delivery on Bone Regeneration Following Composite Bone-Muscle Injury

**DOI:** 10.1101/2020.03.25.008813

**Authors:** Ramesh Subbiah, Albert Cheng, Marissa A. Ruehle, Marian H. Hettiaratchi, Luiz E. Bertassoni, Robert E. Guldberg

## Abstract

The objective of this study was to investigate the controlled release of two growth factors (BMP-2 and VEGF) as a treatment strategy for clinically challenging composite injuries, consisting of a segmental bone defect and volumetric muscle loss. This is the first investigation of dual growth factor delivery in a composite injury model using an injectable smart delivery system consisting of heparin microparticles and alginate gel. The loading efficiency of growth factors into these biomaterials was found to be >90%, revealing a strong affinity of VEGF and BMP-2 to heparin and alginate. The system could achieve simultaneous or sequential release of VEGF and BMP-2 by varying the loading strategy. Single growth factor delivery (VEGF or BMP-2 alone) significantly enhanced vascular growth *in vitro*. However, no synergistic effect was observed for dual growth factor (BMP-2 + VEGF) delivery. Effective bone healing was achieved in all treatment groups (BMP-2, simultaneous or sequential delivery of BMP-2 and VEGF) in the composite injury model. The mechanics of the regenerated bone reached a maximum strength of ∼52% of intact bone with sequential delivery of VEGF and BMP-2. Overall, simultaneous or sequential co-delivery of low-dose BMP-2 and VEGF failed to fully restore the mechanics of bone in this injury model. Given the severity of the composite injury, VEGF alone may not be sufficient to establish mature and stable blood vessels when compared with previous studies co-delivering BMP-2+VEGF enhanced bone tissue regeneration. Hence, future studies are warranted to develop an alternative treatment strategy focusing on better control over growth factor dose, spatiotemporal delivery, and additional growth factors to regenerate fully functional bone tissue.

**Highlights:**

- We developed a smart growth factor delivery system using heparin microparticles and alginate that facilitates tunable delivery of VEGF and BMP-2 in a simultaneous or sequential manner by merely varying the loading strategy.
- *In vitro*, both VEGF and BMP-2 alone promoted vascular growth; however, VEGF was significantly more potent, and there was no detectable benefit of co-delivery.
- *In vivo*, both BMP-2 alone and co-delivery of VEGF and BMP-2 promoted bone formation in the challenging bone/muscle polytrauma model; however, none of the treatment groups restored biomechanical properties to that of uninjured bone.

## 1. Introduction

Severe musculoskeletal injuries are highly prevalent in civilian and military trauma and can cause permanent disability. Polytrauma (type IIIA/B/C open fractures) involves composite injury of multiple tissues, such as a segmental bone defect with volumetric muscle loss (VML). This traumatic condition can often result in complex orthopedic problems such as delayed fracture union or non-union and is frequently associated with patient morbidity and chronic pain.[1-4] There currently are no clinical therapies that regenerate lost muscle,[5] and treatment of chronic non-union and severe composite injury requires multiple surgical procedures that increase the financial, physical, and emotional burden on patients.[3] Current gold standard treatments for non-union open fracture rely on bone autografts, which are costly, invasive, associated with donor-site morbidity, and challenging to scale up in size and number of autografts necessary to treat all patients.[6] Alternatively, acute non-union fractures can be treated by providing both mechanical support, such as fracture stabilization and immobilization, and biological support such as with the use of growth factors.[7]

Growth factors are soluble proteins that actively participate in cell signaling pathways, and regulate cell behavior, including proliferation, self-renewal, migration, and differentiation.[8] Preclinical studies using diverse injury models have focused on understanding the critical role of exogenous growth factor delivery in stimulating regeneration in various organ systems. Bone morphogenetic protein-2 (BMP-2) and BMP-7 are osteoinductive agents that can stimulate significant bone healing in both preclinical models and human patients. Thus, treatment using BMP-2 and BMP-7 has emerged as an alternative to autograft bone tissues for repairing long bone fractures.[8-12] We have developed and tested an alginate-based delivery system for BMP-2 that provides sustained protein delivery to femoral bone defects in rats.[13-15] We have investigated a range of BMP-2 doses using this delivery system (1 μg to 30 μg) and have demonstrated successful bone healing in a dose-dependent manner.[16-18] However, high doses (30 µg) of BMP-2 in rodents, similar to what has been used in the clinic (0.1-1 mg BMP-2/kg body weight), have resulted in side effects, such as heterotopic ossification and inflammation.[18-20] This suggests that an optimized dose and smart delivery system that provides spatiotemporal control over protein delivery is necessary to avoid these side effects while achieving effective bone regeneration.

We have developed a preclinical rodent model to mimic composite bone-muscle injuries in human patients and have demonstrated that a concomitant muscle injury attenuates bone regeneration, even with BMP-2 treatment at the doses that result in significant bone healing in a segmental bone defect model in a rodent (2-2.5 µg).[17, 21] Previous studies demonstrated that inadequate revascularization, dysregulated inflammation, fibrosis, chronic loss of limb function, circulating stem cells, and myokines progressively worsened the capacity for bone healing over time in composite injuries.[4, 21] Restoration of the vascular network at the defect site by delivering angiogenic growth factors may improve the bone healing effect of BMP-2 in the composite injury model. Vascularization in response to tissue injury is mainly governed by thrombin, fibrinogen fragments, and other proangiogenic growth factors. Platelet-derived growth factor (PDGF), vascular endothelial growth factor (VEGF), transforming growth factors (TGF-α, TGF-β), basic fibroblast growth factor (bFGF), platelet-derived endothelial cell growth factor (PD-ECGF), and angiopoietin-1 (Ang1) are several essential angiogenic growth factors that stimulate endothelial proliferation, migration, and tube formation.[8, 22].

The selection and optimization of appropriate growth factors, doses, and an ideal smart delivery system are critical to achieving optimal protein usage and successfully enhancing regeneration of complex tissue injuries such as composite injury. An optimized smart delivery system should demonstrate high protein loading efficiency, easy administration, significantly controlled delivery to targeted tissue, and minimal adverse effects.[8, 11, 23] In this study, we used an injectable smart delivery system consisting of heparin microparticles (HMPs) and alginate gel to deliver VEGF and BMP-2; this system relies on electrostatic interactions between the materials (alginate, HMPs) and growth factors (VEGF, BMP-2) to control growth factor release that mimics the natural healing cascade. This delivery system is capable of delivering multiple growth factors in a simultaneous and sequential manner by varying the loading strategy. This is the first study to test the effects of dual growth factor (low-dose BMP-2 and VEGF) delivery on functional bone regeneration in a critical composite injury model that represents a clinical polytraumatic condition. Single growth factor delivery (VEGF or BMP-2) significantly enhanced vascular growth *in vitro*, but no synergistic effect was observed for BMP-2 and VEGF co-delivery. All treatment groups (BMP-2, simultaneous or sequential delivery of BMP-2 and VEGF) resulted in effective bone healing as measured by bone volume, mineral density, and stiffness of the regenerated bone *in vivo*, although the failure strength of the regenerated bone was not fully restored to that of intact, uninjured bone. Together, these data demonstrate the need to evaluate the molecular and cellular events occurring in this injury model and develop an alternative treatment strategy that not only enables better control over growth factor delivery and dose but also mimics natural bone repair.

## 2. Materials and methods

All animal experiments were performed in accordance with protocols approved by the Georgia Institute of Technology Institutional Animal Care and Use Committee (IACUC).

### 2.1. Smart delivery system

Heparin methacrylamide microparticles (HMPs) were prepared following our previously reported protocol.[24] Briefly, heparin ammonium salt (17–19 kDa; Sigma-Aldrich, St. Louis, MO) was covalently functionalized with N-(3-Aminopropyl)methacrylamide (APMAm; Polysciences, Warrington, PA) using 1-ethyl-3-(3-dimethylaminopropyl)carbodiimide (EDC; Thermoscientific, Rockford, IL) and N-hydroxysulfosuccinimide (Sulfo-NHS; Thermoscientific, Rockford, IL) (**Figure 1A**). The resultant heparin methacrylamide was then dialyzed against water using dialysis tubing with MW cutoff of 3.5 kDa (Spectrum Laboratories, Rancho Dominguez, CA) and then lyophilized (Labconco, Kansas City, MO) for four days. The microparticles of heparin methacrylamide were fabricated using the water-in-oil (W/O) emulsion method. Briefly, heparin methacrylamide was dissolved in phosphate-buffered saline (PBS; Corning Mediatech, Manassas, VA) and gently mixed with equimolar amounts of ammonium persulfate (Sigma Aldrich) and tetramethylethylenediamine (Sigma Aldrich) and slowly added to the mixture of corn oil and polysorbate 20 (Promega, Madison, WI), followed by homogenization at 3000 rpm for 5 minutes using a Polytron PT3100 homogenizer (Kinematica, Switzerland) to create a W/O emulsion. The entire procedure was carried out at a cold temperature using an ice bath. The heparin methacrylamide emulsion was then thermally crosslinked with constant stirring at 55°C using a water bath under inert conditions (N_2_ gas purging) for 30 minutes, then centrifuged at 3000 rpm for at least 10 minutes to remove the oil phase, and then washed and centrifuged with acetone and water respectively to remove excess oil or loosely cross-linked HMPs. The HMPs were disinfected using 70% ethanol solution, then washed with sterile water, lyophilized and stored at 4°C until use. The mean size, distribution, and the polydispersity index (PDI) of HMPs were determined using a Brookhaven 90 Plus Particle Size Analyzer. The prepared HMPs were then stained with 0.1% (w/v) Safranin-O for 30 mins, washed, and observed under a light microscope to confirm the presence of heparin and visualize the particle morphology (Carl Zeiss, Germany). Lyophilized HMPs were resuspended in 70% ethanol and placed on an SEM mounting stud covered in copper tape and left until the ethanol evaporated. HMPs were sputter-coated with gold using a Hummer 5 Gold/Palladium Sputtering (Anatech, Union City, CA) and imaged on an FEI Nova Nanolab 200 Focused Ion Beam/Scanning Electron Microscope (SEM; FEI, Hillsboro, OR). The smart delivery system (HMPs in alginate gel) was prepared as follows. Briefly, RGD-alginate (PRONOVA UP MVG; Viscosity > 200 mPa*s; MW ∼270 kDa; with a G/M ratio of ≤ 1.5) was dissolved in αMEM, mixed with HMPs and CaSO_4_ (Sigma, USA) and stored at 4°C. Two syringes and a disposable three-way stopcock (HS-T-01, Hyupsung Medical Co., Ltd, Korea) were used to thoroughly mix the solutions, as previously reported.[25]

**Figure 1.**
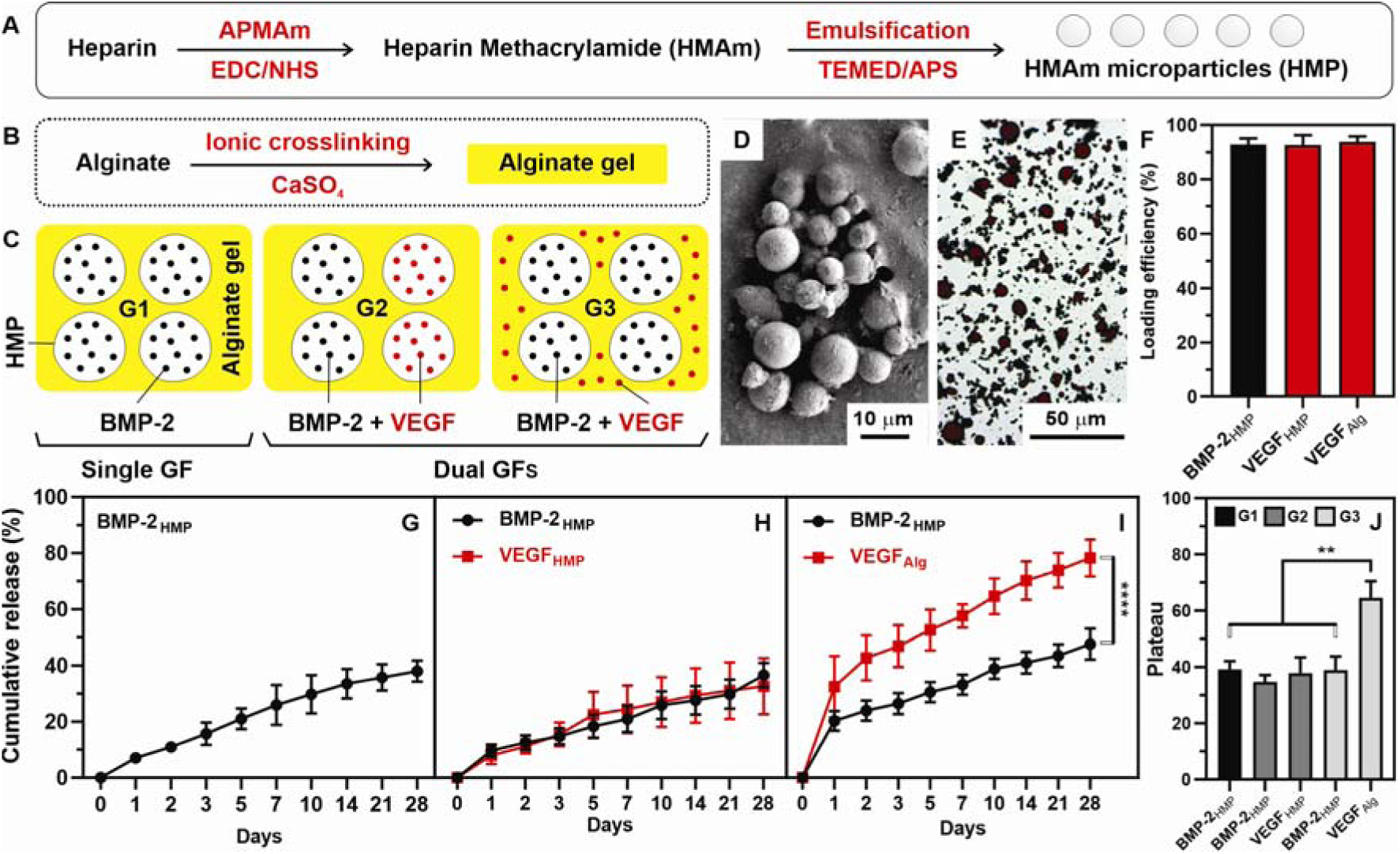
Smart delivery system. A) Schematic depicts the synthesis of heparin methacrylamide (HMAm) and preparation of HMAm microparticles (HMP), and B) fabrication of injectable alginate gel which consists of growth factor loaded-HMP. C) Schematic describes the test groups: G1 = BMP-2-loaded HMPs distributed in alginate gel, G2 = BMP-2 + VEGF loaded HMPs distributed in alginate gel, and G3 = BMP-2 loaded HMPs distributed in VEGF loaded alginate gel. D) Scanning electron microscopy shows the polydisperse and spherical structure of the microparticles (scale bar = 10 μm), and E) Safranin O staining confirms the presence of heparin (scale bar = 50 μm). F) Growth factors loading efficiency was found to be >90% for BMP-2 and VEGF on HMPs and VEGF on alginate. G) Release of BMP-2 from HMP (G), BMP-2/VEGF from HMP (H), and BMP-2 from HMP and VEGF from alginate (I), and nonlinear fit analysis for BMP-2 and VEGF (J), over 28 days in PBS at 4□. Significant differences in growth factors release between alginate and HMPs **** p≤0.0001, ** p ≤ 0.01.

### 2.2. Growth factors loading and release

VEGF165 was purchased from R&D Systems, Minnesota, USA, and BMP-2 was received from Pfizer, Inc., New York City, NY. The loading efficiency of the HMPs and alginate gel for human VEGF (pI = 8.5) and human BMP-2 (pI = 9.0) was investigated using 100 ng/mL of each growth factor. For the loading procedure of G1 and G2, HMPs were incubated with either a single or dual growth factor solution (**Figure 1C: G1, G2**) in 1 mL of 0.1% (w/v) bovine serum albumin (BSA; Millipore Corporation, Billerica, MA) in PBS, and rotated at 4°C overnight. The HMPs were then centrifuged, and the unbound growth factors in the supernatant were measured using enzyme-linked immunosorbent assay (ELISA; R&D Systems, Minneapolis, MN). For the other loading procedure (G3), VEGF was entrapped in alginate and mixed with BMP-2-loaded HMPs during the gel preparation as discussed earlier (**Figure 1C: G3**). The loaded alginate gel was washed with 0.1% BSA and analyzed for unbound growth factor at time 0h using ELISA. Growth factor loading efficiency was determined by subtracting the amount of growth factors remaining in the supernatant or washed solution from HMPs or alginate gel from the initial amount of growth factors used for loading. Passive release of BMP-2 into 0.1% (w/v) BSA in PBS was monitored over a period of 28 days at 4°C. Alginate gel with or without HMPs was centrifuged at 3000 rpm for 5 minutes to obtain released growth factor at various time points over 28 days and replaced with an equivalent volume of fresh 0.1% (w/v) BSA in PBS. Release samples were then analyzed for growth factor content using ELISA.

### 2.3. Effect of VEGF/BMP-2 on microvascular fragments (MVFs) growth

MVFs were isolated from the epididymal fat pads of two retired breeder Lewis rats, as previously described.[26] Fat tissue (10 – 12 ml) was minced, digested with a collagenase solution for 7 min at 37 □C, and tissues between 50 and 500 µm were obtained using selective membrane filters with the pore size of 200 µm and 20 µm to remove partially undigested adipose tissue and single cells, respectively. MVFs were suspended at a density of 20,000 fragments/mL in solution prior to gelation. Collagen gels were made at 3% (w/v), buffered with Dulbecco’s modified eagle medium (DMEM; Thermo-Fisher Scientific; Waltham, MA), and mixed with growth factor-loaded HMPs (G2, G3, G4) as schematically illustrated in **Figure 2A**.[17] The effect of growth factors on MVF growth in the disc-shaped collagen gel was measured. At least three independent gels were evaluated per condition. Briefly, gels were fixed with 4% paraformaldehyde and stained with 10 mg/mL rhodamine-labeled Griffonia simplicifolia lectin I (GS-1lectin; Vector Laboratories; Burlingame, CA). Five randomly selected fields were imaged per gel to an area of 3×3 mm and a depth of ∼500 µm using a Zeiss LSM 700 confocal microscope. Maximum intensity z-projections were created for each stack, thresholded, and skeletonized using the AngioAnalyzer ImageJ plugin to determine branch number and total length.

**Figure 2.**
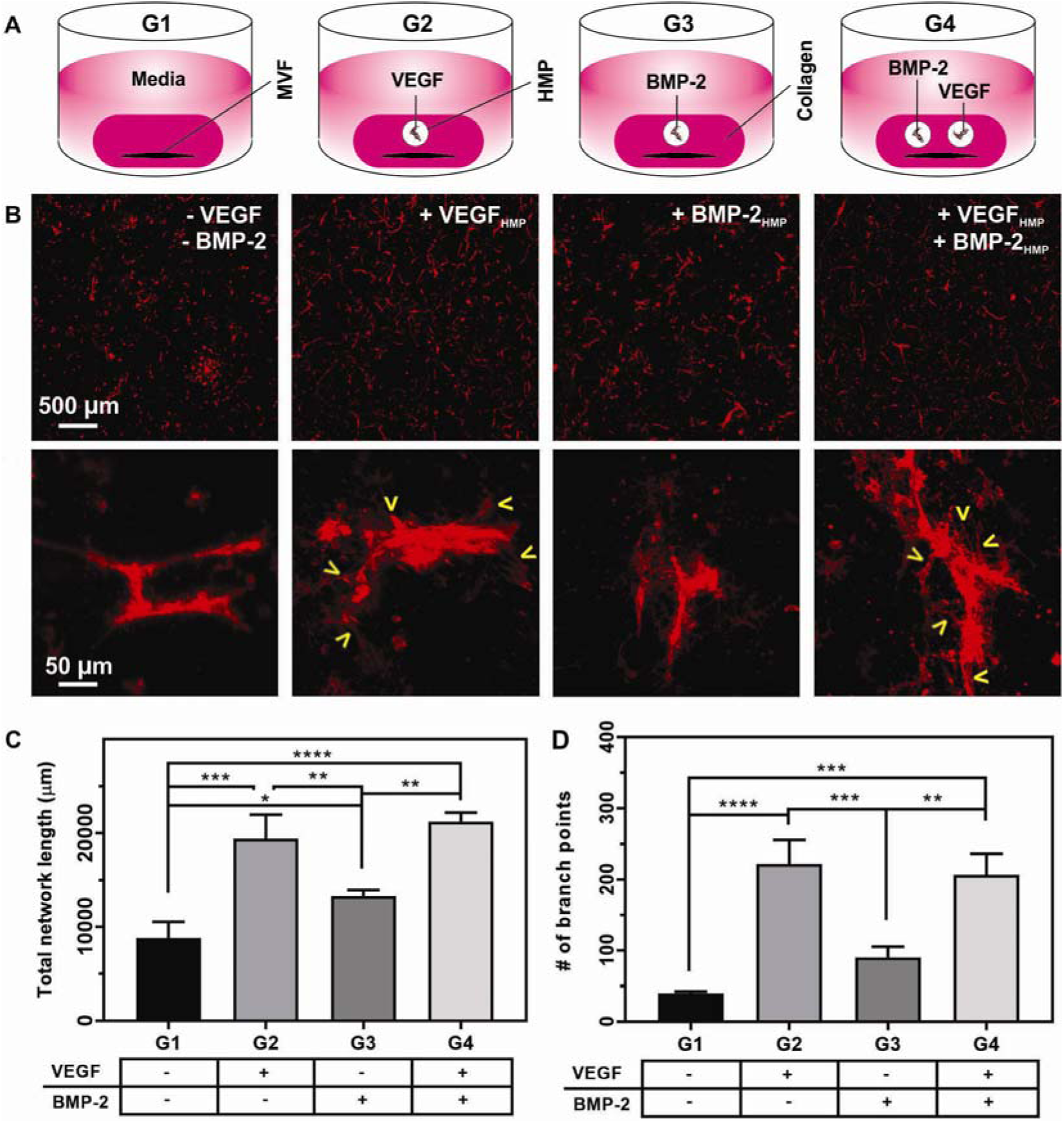
Microvascular network growth analysis *in vitro*. A) Schematic depicts the MVF impregnated collagen gel, supported with no growth factor (G1), VEGF alone (G2), BMP-2 alone (G3), and VEGF+BMP-2 (G4), respectively. Growth factors are delivered in the gel using HMPs. B) Low (top) and high (bottom) magnification z-stack images of MVF in collagen gel constructs at day 10 of culture (red - GS1-lectin-stained MVF), G1, G3 constructs illustrate less network formation than G2 and G4, respectively. C) Quantification of total network lengths and D) Quantification of branch numbers between groups. G2 and G4 exhibit higher total network length and number of branch points than G1 and G3 (mean□±□SD, n□=□3 per group, statistical significance *p□<□0.05, **p<0.005, ***p<0.001, ****<0.0001).

### 2.4. Animals

For these studies, 13-weeks-old female Sprague-Dawley rats were obtained from Charles River Laboratories (Wilmington, MA, USA). Rats were double-housed in individually ventilated caging (Tecniplast, West Chester, PA, USA) with bedding of corn cob, processed paper, and with tunnel and gnawing blocks (Bio-Serv, Prospect, CT, USA) for enrichment. Rats were maintained on a 12-h light/dark cycle and allowed *ad libitum* access to food (Purina Mills International #5001) and filtered tap water treated with ultraviolet light in bottles. All animals were allowed to acclimate for at least 2 weeks before any procedures were performed. After each procedure, a divider was temporarily placed in the cage for better monitoring of postoperative recovery. Animals were randomly assigned to treatment groups.

### 2.5. Surgical procedure

Unilateral composite bone and muscle injuries (**Figure 3B**) were created in 13-weeks-old female Sprague-Dawley rats as previously described.[21] Isoflurane (Henry Schein Animal Health, Dublin, OH, USA) was used to anesthetize the rats. An anterolateral skin incision was then made in the thigh, and overlying muscle was separated by blunt dissection to create a muscle window to reach the femur bone and to place a radiolucent polysulfone fixation plate for internal stabilization. A critically sized 8-mm defect was created in the mid-diaphysis of the femur using an oscillating saw, and a concomitant 8 mm diameter full-thickness quadriceps femoris muscle defect was made. Alginate gels containing growth factors were injected into an electrospun polycaprolactone (PCL) nanofiber mesh tube placed at the composite defect site, as previously described (**Figure 3B**).[15] The treatment groups were G1: 5 µg BMP-2 loaded on HMPs and mixed in alginate gel (controlled release of single growth factor; n=4); G2: 5 µg of BMP-2 and 5 µg of VEGF loaded on HMPs and mixed in alginate gel (simultaneous release of dual growth factor; n=8); and G3: 5 µg of BMP-2 loaded on HMPs and 5 µg of VEGF entrapped in alginate gel (sequential release of dual growth factor; n=4). Finally, the muscle and skin were closed using 4-0 vicryl suture and wound clips, respectively. Before surgery, all animals were given a subcutaneous injection of sustained-release buprenorphine (ZooPharm, Windsor, CO, USA) for analgesia. At 12 weeks post-surgery, animals were euthanized by CO_2_ inhalation.

**Figure 3.**
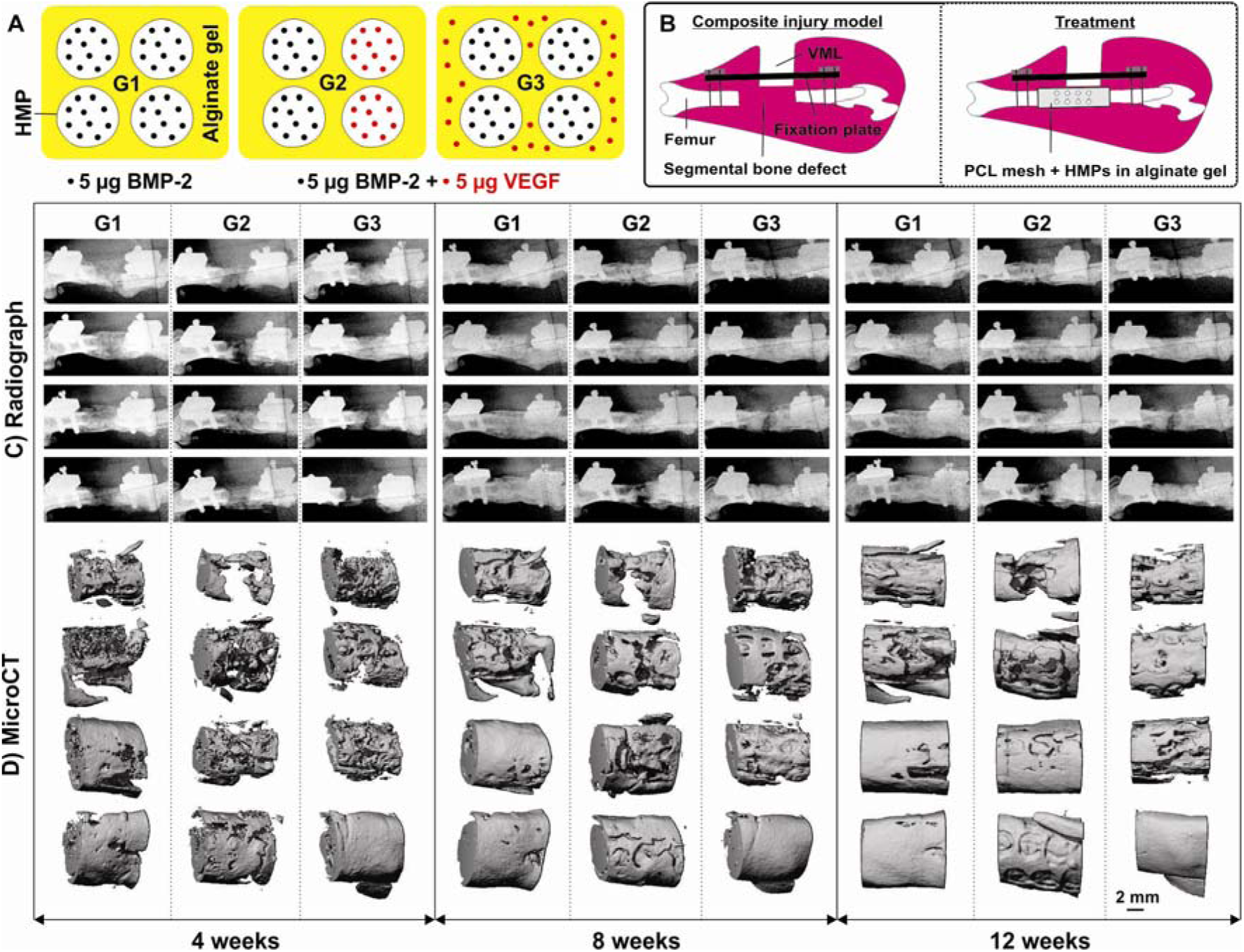
Qualitative analysis of the regenerated bone in composite injury. A) Schematic depicts the HMPs/alginate gel-based delivery systems, the dose of single and dual GF, respectively. B) Schematic shows the composite injury model, which consists of ∼8 mm sized segmental bone defect with ∼280 mg sized volumetric muscle loss (VML) and the treatment using a cylindrical PCL mesh scaffold. C) Longitudinal radiographs and D) longitudinal microCT reconstructed images of each treatment group at weeks 4, 8, and 12.

### 2.6. Bone regeneration analysis

Bone regeneration was assessed longitudinally by radiography and micro-computed tomography (µCT) 4, 8, and 12-weeks post-surgery. The defect region was scanned with a voxel size of 38.9 mm, and the bone volume was quantified by Scanco software using a threshold corresponding to 50% of intact cortical bone.[16, 21] Mechanical testing and histology were performed at the 12-week endpoint after euthanasia. Thighs were harvested, soft tissue was cleared, and fixation plates were removed. Femur ends were potted in Wood’s metal (AlfaAesar) and tested in torsion at a rate of 3 □/s to failure (ELF3200, TA ElectroForce).[13] Maximum torque (torque at failure) and torsional stiffness (linear region of torque vs. rotation) were calculated for all samples. Following mechanical testing, the samples were fixed in 10% neutral buffered formalin (NBF) for 48 hours at room temperature. Samples were then switched to PBS and sent to HistoTox Labs (Boulder, CO, USA) for decalcification, trimming, sectioning (5 µm thickness), and staining with hematoxylin and eosin (H&E) or Safranin□O and Fast Green. Masson’s Trichrome was performed in□house according to established protocols. Bone sections were also stained with 10□μg/ml of GS-1lectin and counterstained with DAPI (ThermoFisher) diluted 1:1,000 in PBS to identify blood vessels. All histology images were taken in the center of the bone defect at 20x magnification using an inverted fluorescence microscope (FL Auto, Evos).

### 2.7. Statistical analysis

Quantitative data were analyzed using GraphPad Prism 7 and expressed as mean ± standard error of the mean (SEM). Statistical significance was determined using two-way ANOVA with Tukey’s post-hoc test.

## 3. Results and discussion

### 3.1. Smart delivery system enables tunable delivery of multiple growth factors

Composite bone-muscle injuries are hard to treat due to muscle loss, inadequate vascularization at the defect site, dysregulated inflammation, and fibrosis.[4, 21, 27] There are currently no regenerative standards of care for volumetric muscle loss which is a common clinical outcome after polytrauma.[5] Although bone autografts are considered gold standard treatments for delayed union and non-union of fractures, they are not ideal due to limited availability of graft, donor site morbidity, increased surgical time, and blood loss.[6]

Tissue healing is regulated by the coordinated release of multiple growth factors by cells with spatiotemporal control. Typically, VEGF is expressed in the early stage and BMP-2 is expressed in the later stage of the natural bone healing process.[28-31] Controlled delivery of osteogenic growth factors such as BMP-2 using a collagen sponge delivery system has provided an alternative treatment strategy to autografts and allografts in the treatment of critical size non-union bone defects; however, the collagen sponge cannot provide adequate control over BMP-2 delivery *in vivo*, resulting in rapid protein release from the injury site.[8, 16] BMP-2 at high doses can elicit side effects such as inflammation and heterotopic ossification;[18-20] thus, reducing the therapeutic dose of BMP-2 is recommended. Additionally, low-dose BMP-2 alone cannot heal more complex injuries such as composite bone-muscle injury.[21] This highlights the need to develop an alternative treatment strategy.

The delivery of multiple growth factors may be advantageous for facilitating complex tissue repair compared to the delivery of a single growth factor and can minimize the dose required for therapeutic effects. Previous studies reported that co-delivery of VEGF with BMP-2 enhanced the bone healing effect of BMP-2 by interacting with BMP-2 and regulating its biological activity.[25, 32-35]In particular, sequential delivery of VEGF followed by BMP-2 improved bone healing compared to simultaneous VEGF and BMP-2 delivery.[25, 35] In contrast, other studies demonstrated insignificant effects of VEGF and BMP-2 on bone healing.[36, 37] Challenges with growth factor efficacy *in vivo* precipitate the need for better biomaterial delivery vehicles that achieve more efficient growth factor loading, preserve protein bioactivity, and improve protein pharmacokinetics and pharmacodynamics.[8]

Achieving physiologically relevant control over growth factor release *in vivo* is challenging using conventional delivery systems. Here, we employed a biomaterials-based smart delivery system consisting of HMPs in alginate gel to closely mimic the physiological growth factor release cascade of bone tissue healing.[31] Heparin is a highly sulfated glycosaminoglycan, and alginate is a polysaccharide polymer. These anionic polymers mimic the extracellular matrix, exhibit electrostatic affinity to cationic growth factors, preserve the biological activity of proteins, and facilitate sustained delivery of growth factors at the injury site while avoiding covalent binding and structural changes of growth factors.[8, 24] We have previously demonstrated that HMPs can be used to effectively control high doses of BMP-2 delivered within a femoral bone defect, resulting in better spatial localization of growth factor delivery.[38] In contrast, BMP-2 at low doses strongly bound to HMPs, resulting in attenuated BMP-2 release and subsequent bone healing in femoral defects.[39] Co-delivery of BMP-2 and VEGF is believed to augment the delivery of low-dose BMP-2. Since heparin can bind a number of cationic growth factors including BMP-2 and VEGF, HMPs were expected to be an ideal delivery vehicle for multiple growth factor delivery.

HMPs were prepared from methacrylamide-functionalized heparin by emulsification methods, following our previous study (**Figure 1A)**.[24] HMPs were spherically shaped as confirmed by optical microscopic and SEM images (**Figure 1 D, E**). The average size of the HMPs was 5.9±1.6 µm with a polydispersity index (PDI) of 0.45. The larger observed PDI is due to the agglomeration of HMPs. HMPs in this study were used to deliver BMP-2 alone (G1) and to co-deliver BMP-2 with VEGF (G2). The bound growth factor fraction was above 90% for both BMP-2 and VEGF when loaded alone or simultaneously, verifying strong binding affinity of the cationic growth factors to anionic HMPs (**Figure 1F**). HMPs minimized the burst release of the growth factors and effectively controlled the release of BMP-2 (∼36.6%) and VEGF (∼32.6%) in a temporal and sustainable manner over a period of 28 days (**Figure 1G-I**). The cumulative release of growth factors from HMPs was found to be between 30% and 40% in all conditions. HMPs demonstrated high loading efficiency and were capable of delivering multiple growth factors simultaneously. We expect that HMPs were also able to protect growth factors from premature degradation, based on previous results.[39, 40]

To provide sequential release of VEGF, followed by BMP-2, VEGF was directly loaded into a second delivery vehicle (alginate), as schematically illustrated in **Figure 1C, I**. Alginate is another popular naturally-derived biomaterial used in tissue engineering applications, offering biocompatibility, biodegradability, and injectability. Alginate is easy to fabricate and entrap cells and growth factors using simple ionic crosslinking (**Figure 1B**).[41-43] Alginate has an isoelectric point of ∼4.95, making it anionic at physiological pH, and exhibits a charge-based affinity towards cationic growth factors, such as VEGF and BMP-2. Herein, VEGF is directly loaded in the alginate gel by physical entrapment (G3), which allows the maximum retention of growth factors and yielded >90% loading efficiency. The cumulative release of VEGF is significantly different when growth factors were loaded in the alginate or HMPs separately (**Figure 1I**). Encapsulation of VEGF in alginate resulted in an early release of VEGF (∼46%) by day 7 and additional sustained growth factor release (∼16.5%) by day 28 while maintaining the controlled release of BMP-2 (∼38.1%) from HMPs in the smart delivery system (**Figure 1I**). This indicates that the cumulative release of BMP-2 and VEGF could be altered by changing the loading strategies (**Figure 1G-J**). Importantly, loading VEGF into the alginate significantly enhanced the early release of VEGF, while loading growth factors onto the HMPs minimized early growth factor release (**Figure 1J**), due to the higher affinity of heparin for cationic growth factors.[24, 38, 40] Overall, we demonstrated that this smart delivery system is versatile since it is capable of delivering multiple growth factors in a tunable and controlled manner simply by electing the material component (HMPs or alginate) to be loaded with the growth factors of interest.

### 3.2. Single growth factor delivery from smart delivery system improves vascularization in vitro

A successful bone regeneration largely depends on adequate vascularization. Prior to *in vivo* studies, the vascularization effects of VEGF and BMP-2 were tested using MVF *in vitro*. Briefly, MVFs were cultured for 9-10 days in three-dimensional collagen gels impregnated with HMPs contains no growth factors (G1), VEGF alone (G2), BMP-2 alone (G3), and both VEGF and BMP-2 (G4) (**Figure 2A**). MVF growth was influenced significantly by VEGF (G2), BMP-2 (G3), and combined VEGF+BMP-2 (G4) as confirmed by the total vascular network length and branch points (**Figure 2B-D**). BMP-2 alone (G3) stimulated MVF network formation, but not to the same extent as VEGF delivery (G2, G4, **Figure 2B**). The observed moderate stimulation of MVF growth by BMP-2 might be due to endothelial cell stimulation and paracrine signalling.[44] MVF growth was not different between VEGF alone (G2) and VEGF+BMP-2 (G4) groups, indicating that there was no additive effect of BMP-2 on vascularization stimulated by VEGF.

### 3.3. Smart delivery system facilitates bone regeneration in a composite bone-muscle injury

The composite injury model used herein consists of an acute femoral bone defect with VML (**Figure 3B**). This rat preclinical model mimics a typical composite muscle-bone injury in humans given that fractures lacking adequate coverage with intact muscle are associated with higher rates of complications and nonunion. We have previously demonstrated that the additional muscle trauma attenuates bone healing at a minimally bridging dose (2.5 µg) of BMP-2 in the segmental bone defect.[17, 21] Given the critical dependence of bone regeneration on adequate vascularization, we hypothesized that effective vascularization would facilitate fracture healing in composite injury. Furthermore, in previous studies, the early release of VEGF enhanced revascularization in both ectopic and orthotopic sites.[25, 35, 37] Thus far, no studies have reported the co-delivery of BMP-2 with VEGF in the composite injury model. This study was hence designed to investigate a smart delivery system to provide sustained release of BMP-2 and VEGF and evaluate the effect of these growth factors on bone regeneration in a composite bone-muscle injury model (**Figure 3A, B**). This is the first investigation of BMP-2 and VEGF co-delivery in this injury model. Three groups were used: BMP-2 (G1), simultaneous delivery (G2), and sequential delivery (G3) of BMP-2 and VEGF. A comparatively low dose (5 µg) of BMP-2 was used to test bone healing in this study, which is much lower than the current FDA approved therapeutic dose (1.5 mg/mL for human use).[19] We have used 5 µg of VEGF loaded into HMPs (G2) or alginate (G3) as a proposed therapeutic approach to enhance revascularization *in vivo* as an alternative to cell-based (MVF) therapy. Radiographs and microCT images demonstrated that defects treated with 5 µg BMP-2 with or without the addition of VEGF showed effective bone regeneration by 12 weeks, resulting in bony bridging across the defect (G1, G2, G3; **Figure 3C, D**). Bone volume and bone mineral density (BMD) in the defect region were measured longitudinally by microCT and found to be comparable for all groups at each time point (**Figure 4A, B**). The torque to failure was found to be comparable for all the groups but did not match that of intact bone (**Figure 4C**). However, the strength of regenerated bone treated by sequential delivery of VEGF then BMP-2 (G3) reached 52.4% of intact contralateral strength when compared to BMP-2 alone (G1, 37.9%), and simultaneous delivery of VEGF and BMP-2 (G2, 37.3%), Likewise, the stiffness of the regenerated bone for all the groups was found to be comparable with the intact contralateral control (**Figure 4D**). Results indicated that there were no significant effects of the added VEGF on bone volume or density irrespective of simultaneous (G2) or sequential (G3) VEGF release kinetics.

**Figure 4.**
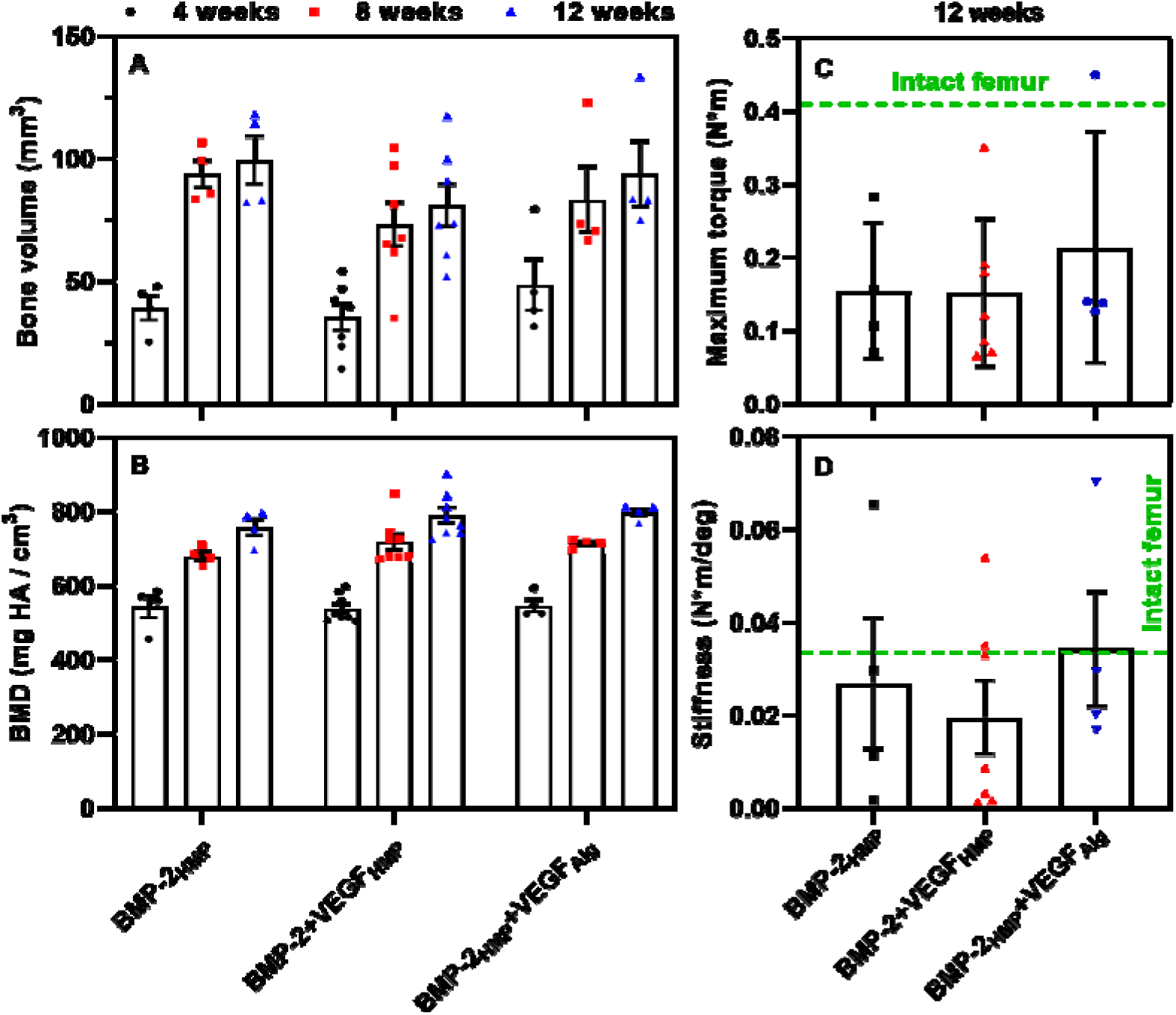
Quantitative analysis of the regenerated bone in composite injury. Longitudinal bone volume (A) and bone mineral density (B) of regenerated bone measured by microCT at 4, 8, and 12 weeks of treatment group delivering BMP-2 by HMPs (G1), BMP-2+VEGF by HMPs (G2), and BMP-2 by HMPs and VEGF by alginate (G3). C) Failure strength and D) stiffness of regenerated bone at 12 weeks, wherein dotted lines indicate the value of intact contralateral femur.

### 3.4. Dual growth factor delivery improves the regenerated bone morphology

Representative samples (n=2) from each treatment group were sectioned and stained with H&E, Saf-O, and Masson’s trichrome. H&E staining (**Figure 5A**) revealed morphologic differences in the tissue formed within the bone defect, particularly between the single and dual growth factor treatment. Areas of new bone formation were observed in all samples, but these areas were much larger and more widespread throughout the defect in the sequentially delivered VEGF (G3) samples. Moreover, groups treated with the early release of VEGF (G3) exhibited the formation of marrow-like structures directly adjacent to these areas of new bone formation (**Figure 5A**). In contrast, the sample treated with BMP-2 alone (G1) demonstrated only small sparse islands of new bone formation surrounded by mostly fibrous tissue. Saf-O staining (**Figure 5B**) revealed negligible cartilage present in the defect at 12 weeks post-surgery for all groups. The presence of alginate and HMPs was observed across the defect site (**Figure 5A, B**). Finally, trichrome staining (**Figure 6A**) demonstrated that the G3 treated group had more areas of collagen fiber alignment in the areas of new bone formation, indicative of mature lamellar bone. Conversely, the BMP-2 alone treated groups (G1) exhibited a much more disorganized collagen structure, typical of woven bone deposition. Overall, the tissue of the defect area showed woven bone tissue in all three groups. However, more mineralized bone tissue and blood vessels around alginate regions were observed at the whole circumference of the defect site in the early release of VEGF (G3) compared to the simultaneous release of VEFG (G2) and BMP-2 alone (G1). Immunofluorescence staining of GS-1 lectin demonstrated that the VEGF delivery enhanced the vascularization in the defect site (**Figure 6B**). All of the treatment groups promoted bone formation, suggesting that our smart delivery system could achieve favourable delivery of growth factors for tissue regeneration. However, none of the tested groups restored biomechanical functions of the native bone in this challenging bone/muscle polytrauma model. This may be because 1) the molecular and cellular events involved in the osteogenic and angiogenic process are critically altered in composite injuries and 2) strong binding of low-dose growth factors with HMPs that attenuates growth factor release. Given the severity of the composite injury, VEGF alone may not be sufficient to establish mature and stable blood vessels when compared with the previous studies in which delivering VEGF enhanced bone tissue regeneration.[25, 32-35] Hence, a future study is warranted to develop alternative strategies including but not limited to optimization of dose, delivery vehicle, additional angiogenic growth factors to augment the regeneration of bone in such severe trauma conditions.

**Figure 5.**
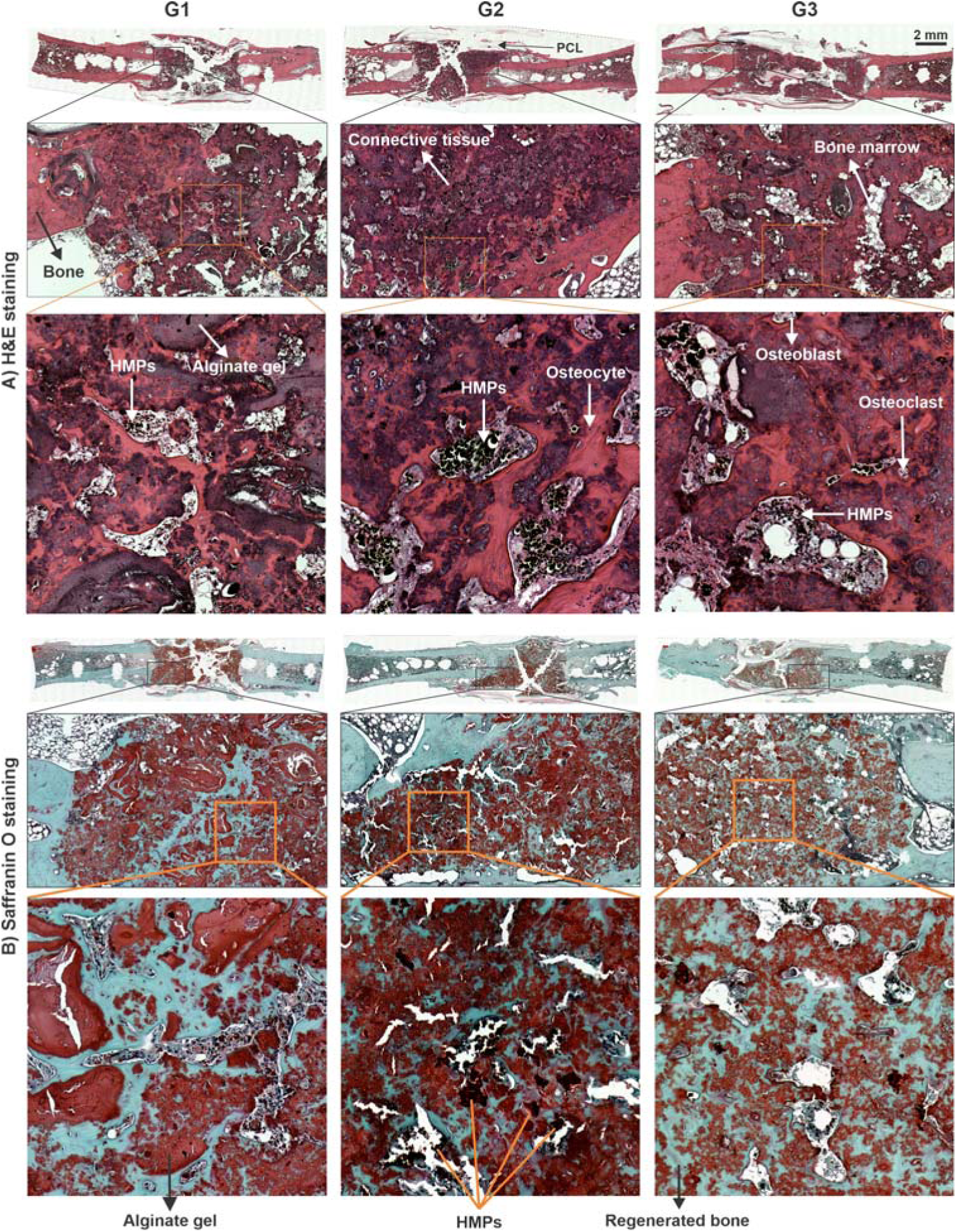
Histology of the segmental defect after 12-weeks treatment (original magnification: 20x). A) H&E staining, bone is presented as a compact structure in a dark pink color, connective tissue in light pink, alginate gel region (isolated pockets) in glossy purple, HMPs in dark red (spherical) and the perforated PCL scaffold structure in white. Areas of direct ossification present structures of firm connective tissue showing positioned collagen fibers. Osteoblast, osteoclast, and osteocytes are denoted in arrows. Mineralized bone tissue was observed at the entire circumference of the defect site. The tissue of the defect area showed a much higher amount of mineralized bone tissue for G3 than G1 and G2. B) Safranin-O staining showed pockets of endochondral ossification (bluish-green), isolated pockets of alginate (glossy pink), spherical HMPs (dark red), and bone marrow structure at the defect site.

**Figure 6.**
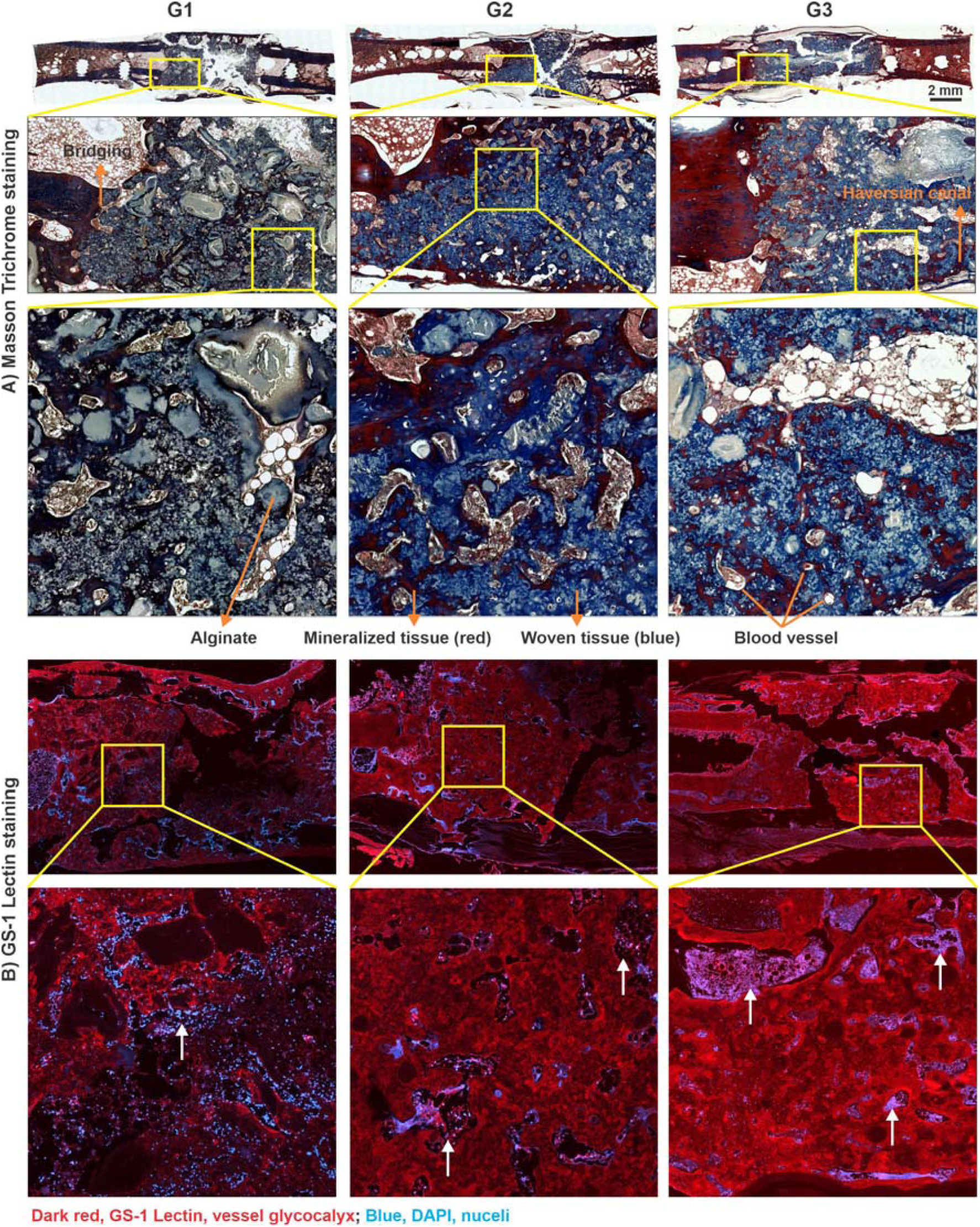
Vascularized bone tissue. A) Masson’s trichrome staining shows a detailed description of new bone formation within the PCL scaffold. Identical images present the Haversian canal, connective tissue, blood vessels, mineralized (red), and general woven bone tissue (blue). B) GS□1 lectin staining for blood vessels within regenerate bone at 12 weeks from each treatment group, and white arrows depict marrow-like tissue mineralized bone.

## 4. Conclusions

The purpose of this study was to evaluate the effects of temporally-controlled dual angiogenic and osteogenic growth factor delivery approaches on vascular growth *in vitro* and bone regeneration in a composite injury model. This smart delivery system (HMPs in alginate gel) exhibited high growth factor binding efficiency and effective delivery of cationic growth factors in a controlled manner while preserving the bioactivity of the growth factors. The *in vitro* study results suggest that the delivery of a single growth factor (VEGF or BMP-2) significantly enhances the growth of MVFs over the control group; however, no synergistic effect was observed for a dual growth factor delivery (BMP-2 + VEGF). The smart delivery system used for the *in vivo* study offered an effective strategy to deliver multiple growth factors in a temporally controlled manner that mimics the physiologically relevant sequential release profile of VEGF followed by BMP-2. This was the first study to test dual BMP-2 and VEGF delivery on the functional regeneration of bone in a challenging preclinical model of established bone non-union defect and muscle injury. *In vivo* results (x-ray and microCT) suggest substantial bone healing in all test groups; however, the mechanics of the regenerated bone were not restored when compared to uninjured bone. Overall, *in vivo* results suggest that VEGF alone was not sufficient to augment the bone healing effect of low-dose BMP-2 in a critical composite injury. Hence, a future study is warranted to develop an alternative treatment strategy.

## Declaration of Competing Interest

The authors declare no competing financial interest.

## Acknowledgments

This work was supported under the AFIRM II (U.S. Armed Forces Institute of Regenerative Medicine) effort, award no. W81XWH-14-2-0003. The U.S. Army Medical Research Acquisition Activity is the awarding and administering acquisition office. Opinions, interpretations, conclusions, and recommendations are those of the authors and are not necessarily endorsed by the Department of Defense. BMP-2 used in this study was provided by Pfizer Inc. The authors wish to acknowledge the core facilities at the Parker H. Petit Institute for Bioengineering and Bioscience at the Georgia Institute of Technology for the use of their shared equipment, services, and expertise. Authors also thank Ryan Akman, Gilad Doron, Brett Klosterhoff, Lina Mancipe-Castro, Casey Vantucci, and Hazel Stevens for their assistance with surgeries. The authors acknowledge Lauren L. Kronebusch for help with manuscript editing.

## Authors’ roles

Conception and study design: RS and REG. Study conduct: RS, AC, MAR, and REG. Data collection, analysis, interpretation, and drafting: RS. Revising manuscript content: MHH, LEB, and REG. Approving final version of manuscript: all authors.

